# N-BLR, a primate-specific non-coding transcript, modulates the epithelial-to-mesenchymal transition and leads to colorectal cancer invasion and migration

**DOI:** 10.1101/004796

**Authors:** Isidore Rigoutsos, Sang Kil Lee, Su Youn Nam, Tina Catela Ivkovic, Martin Pichler, Simona Rossi, Peter Clark, Hui Ling, Yi Jing, Masayoshi Shimizu, Roxana S. Redis, Maitri Y. Shah, Xinna Zhang, Eun Jung Jung, Aristotelis Tsirigos, Li Huang, Jana Ferdin, Roberta Gafà, Riccardo Spizzo, Milena S. Nicoloso, Maryam Shariati, Aida Tiron, Jen Jen Yeh, Raul Teruel, Lianchun Xiao, Sonia A. Melo, Elsa Flores, Massimo Negrini, Menashe Bar-Eli, Sendurai A. Mani, Chang Gong Liu, Ioana Berindan-Neagoe, Manel Esteller, Michael J. Keating, Giovanni Lanza, George A. Calin

**Author notes:** These authors contributed equally (Isidore Rigoutsos, George A. Calin). These authors contributed equally (Isidore Rigoutsos, Sang Kil Lee, Su Youn Nam).

## Abstract

Non-coding RNAs have been commanding increasingly greater attention in recent years as the few that have been functionalized to date play important roles in key cellular processes. Here we show that N-BLR, a ∼900 bp non-coding RNA, modulates the epithelial-to-mesenchymal transition, increases colorectal cancer invasion, and functions as a migration enabler by affecting the expression of ZEB1 and E-cadherin. In patients with colorectal cancer, N-BLR expression associates with tumor stage and invasion potential. As N-BLR contains several instances of a category of DNA motifs known as pyknons, we also designed a custom-made array to investigate the possibility that other pyknon loci may be transcribed. For several of the loci probed by the array we found that the corresponding pyknons are differentially expressed between cancer and normal tissue samples. Taken together the data suggest that a systematic study of other pyknon-containing non-coding RNAs like N-BLR may be warranted in the context of colorectal cancer.

---

Novel experimental methods and recent technological advances have established that, in addition to the protein coding regions, significant parts of the human and other genomes give rise to ncRNAs (Bertone et al. 2004; Carninci et al. 2005; Cheng et al. 2005; Khaitovich et al. 2006). In terms of diversity, ncRNAs handily outnumber protein coding transcripts complicating functional investigations (Louro et al. 2009). Indeed, many classes of *experimentally* identified ncRNAs have been reported in the literature including miRNAs (Bartel 2004), piRNAs (Kim 2006), lincRNAs (Guttman et al. 2009), tiRNAs (Taft et al. 2009), moRNAs (Langenberger et al. 2009), sdRNAs (Taft et al. 2009), or long enhancer ncRNAs (Orom et al. 2010), and others. However, the full complement of ncRNAs and their functional involvement in the regulation of cellular processes and, by extension, in the onset and progression of human disorders continues to remain largely unknown (Mattick 2009).

Among ncRNAs, the best-studied group is that of microRNAs (miRNAs). MiRNAs are short RNAs, 19-23 nucleotides (nts) in length that bind their target mRNAs in a sequence-dependent manner thereby regulating the target's expression (Ambros 2008; Bartel 2009; Rigoutsos 2009). During the last decade, miRNAs have been found to be implicated in many disease settings (Mendell 2008; Vasilescu et al. 2009; Williams et al. 2009; Small and Olson 2011) whereas more recently they were also shown to act as mediators of molecular interactions that obviate direct molecular contact (Poliseno et al. 2010). In the latter context, co-expressed transcripts (protein-coding as well as non-coding) that are targeted by the same collection of miRNAs “compete” for a finite amount of available miRNA molecules (*ceRNAs*) (Poliseno et al. 2010; Karreth et al. 2011; Tay et al. 2011). Changes in the abundance of any one ceRNA can affect those among the remaining ceRNAs that have expression near an underlying transition threshold (Poliseno et al. 2010; Cesana et al. 2011; Mukherji et al. 2011; Sumazin et al. 2011; Tay et al. 2011; Ala et al. 2013).

Long non-coding RNA (lncRNAs) burst onto the scene a few years ago and many are currently known in the public domain (Guttman et al. 2009; Djebali et al. 2012). Even though the full spectrum of ncRNAs remains unclear (Mattick and Makunin 2006) several lncRNAs have been shown to be important in diverse contexts such as chromatin modification and remodeling (Gupta et al. 2010a; Kogo et al. 2011), X chromosome inactivation (Lee and Lu 1999; Ogawa et al. 2008; Zhao et al. 2008; Tian et al. 2010), lineage-specific transcriptional silencing (Pandey et al. 2008), regulation of mRNA export (Chen and Carmichael 2009), activation of a growth-control gene program (Hutchinson et al. 2007; Yang et al. 2011) or of homeobox genes (Wang et al. 2011), lineage-specific silencing (Pandey et al. 2008; Latos et al. 2012), somatic tissue differentiation (Flynn et al. 2012), etc. LncRNAs have also been linked to human conditions such as brachydactyly (Maass et al. 2012) and the Prader-Willi syndrome (Bischof et al. 2007), and to cancers such as melanoma (Khaitan et al. 2011), colon (Calin et al. 2007a; Kogo et al. 2011) and prostate cancer (Prensner et al. 2011).

Pyknons (‘peak-non-s’) are a class of DNA motifs that were initially identified computationally using an *unsupervised* pattern discovery process (Rigoutsos and Floratos 1998; Rigoutsos et al. 2006). The motifs are ≥ 16 nts long and have ≥ 30 exact copies in the *intergenic* and *intronic* space of the genome; what makes these motifs intriguing is that nearly all messenger RNAs (mRNAs) contain one or more pyknons in them in orientation that is sense to the mRNA suggesting the possibility that mRNAs and ncRNAs sharing the same pyknons act as ceRNAs for one another. Pyknon sequences are organism-specific: in fact, across human and mouse, we have found that the respective sets of pyknon motifs are neither syntenic nor conserved at the sequence level (Rigoutsos et al. 2006; Tsirigos and Rigoutsos 2008). Despite this absence of synteny and *<u>sequence</u>* conservation, as we showed previously, the intronic instances of human and mouse pyknons capture widespread *<u>functional</u>* conservation (Tsirigos and Rigoutsos 2008). Virtually all pyknon sequences are *sense* to the spliced mRNAs of some protein-coding genes and *antisense* to the introns of other protein-coding genes: these sense/antisense relationships between mRNAs and introns suggest intriguing possibilities for *trans* regulatory control (Tsirigos and Rigoutsos 2008) a possibility that received experimental support (Robine et al. 2009; Saito et al. 2009; Rigoutsos 2010). Pyknons have also been reported in plants where they are found to have the same properties as their animal counterparts (Feng et al. 2009). More recently, it was reported that the DNA methyltransferase DNMT1 binds RNA at pyknon loci and that the corresponding regions are hypomethylated (Di Ruscio et al. 2013).

Here we describe our discovery and characterization of N-BLR (pronounced: e**N**a**BL**e**R**), a pyknon-containing lncRNA. We examine N-BLR's expression between normal colon and colorectal cancer (CRC) and describe its role in <u>enabling</u> invasion and migration by modulating the epithelial-to-mesenchymal transition (EMT). The organism-specific aspect of the pyknon motifs, the realization that some of the corresponding genomic loci are transcribed into RNAs, and the fact that a pyknon-containing transcript (N-BLR) is intimately linked to disease, prompted us to explore further the possibility of transcription at other pyknon loci as well as to investigate the possibility of a connection to disease that extends beyond N-BLR. To this end, we designed a custom microarray aimed at probing the genome at more than 1,000 pyknon loci and used it to examine evidence of transcription at those loci for several tissues, and for both normal and disease samples. It is important to emphasize that working with sequences which repeat multiple times in the human genome has its own inherent complications. We remained mindful of this fact throughout the duration of our project: in the case of the used primers, we ensured that they are unique; in the case of the probes that we designed for our array, we intentionally mixed probes that are unique to the genome with probes that are not and discuss the rationale for this choice in more detail in the Results and Discussion sections.

## Results

### Transcription of genomic regions containing pyknons correlates with clinical parameters and overall survival in colorectal cancer

When we embarked on this project it was not clear whether pyknons capture “passive” DNA motifs (e.g. genomic locations to which molecules bound) or “active” sources of novel transcripts. Thus, we sought to examine the possibility of transcription by designing qRT-PCR assays for 11 genomic pyknon instances selected within regions associated with loss of heterozygozity (LOH) or within known fragile sites (Calin et al. 2004). In what follows, we denote these regions of interest as pyk-reg-14, pyk-reg-17, pyk-reg-26, pyk-reg-27, pyk-reg-40, pyk-reg-41, pyk-reg-42, pyk-reg-43, pyk-reg-44, pyk-reg-83, and pyk-reg-90 (**Supp. Table 1** and **Supp. Table 2**). Owing to our long-standing interest in colorectal cancer (CRC), we used the 11 assays to explore the possibility of transcription across the following seven cell lines: Colo320, SW480, HCT116, IS174, HT-29, Colo205, and SW620. We observed transcription from all 11 genomic pyknon locations with expression levels that varied among the seven cell lines (**Supp. Fig. 1**).

These initial findings spurred us to focus our investigations on tissue samples from human normal colon and CRC. We used qRT-PCR to evaluate the 11 regions of interest in a set of 82 tumor (“sample set A” – **Supp. Table 3A**) and 28 adjacent normal mucosa samples of Caucasian ancestry. In this set of samples, we found significant expression differences for six of the 11 loci in cancer compared to normal tissue: pyk-reg-14, pyk-reg-40, pyk-reg-41, pyk-reg-42, pyk-reg-44, and pyk-reg-90 (Fig. 1a). Additionally, for five loci we detected significant differences between microsatellite stable (MSS) and instable (MSI) CRCs (Fig. 1b). One of the loci in particular, pyk-reg-90, stood apart from the rest: multivariate logistic regression analysis revealed a significant correlation between high expression of pykreg-90 and high tumor stage. The result was arrived at by analyzing 127 patient samples (we combined sample sets A and B in order to reach the higher number of patients needed for the statistical analysis) and had an associated odds ratio=3.682, *p*=0.010 (see **Supp. Table 4**). Moreover, we found that high pyk-reg-90 expression was also associated with poor overall survival in the combined cohort (sample sets A and B) of 127 patients (*p*=0.016, Fig. 1c). We examined a third independent set (“sample set C”) consisting exclusively of 21 metastatic CRC patient-derived <u>xenografts</u> and found pyk-reg-90 to be highly expressed compared to U6 and to be present in 15 of the 21 samples (*p*=0.0258 when compared to the probability of observing this frequency accidentally, **Supp. Figure 2**).

**Figure 1.**
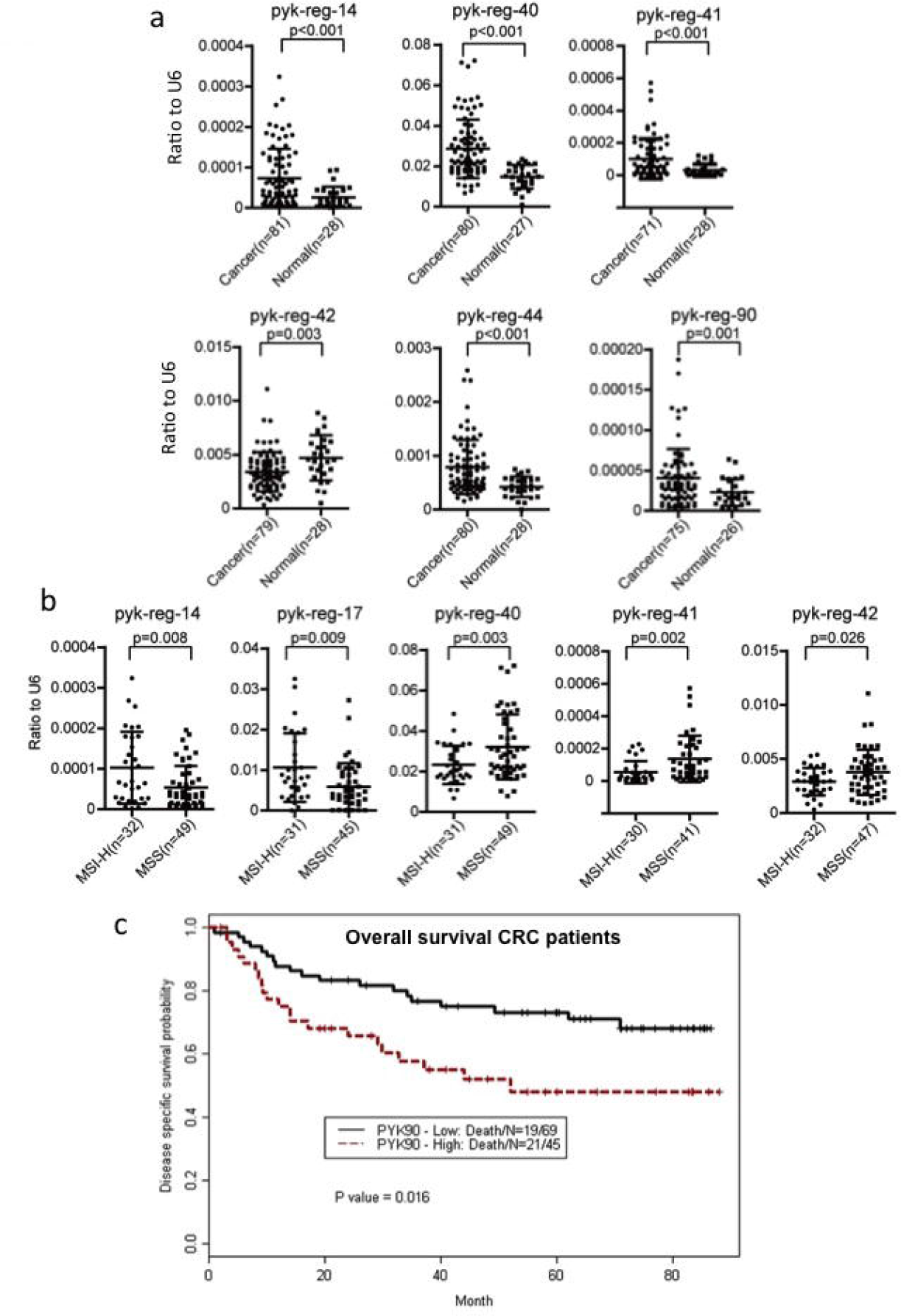
Pyknon locus expression in CRC samples by qRT-PCR. **a,** Expression and distribution of pyknon-regions were analyzed between CRC and paired normal samples (set A – see text) by qRT-PCR. **b,** Expression and distribution of pyknon-regions were analyzed between microsatellite stable (MSS) and microsatellite instable (MSI-H) colorectal cancer by qRT-PCR. The number of samples with measurable expression values (under Ct of 35) is presented in parentheses. The numbers of cancer and normal samples in some cases differ from one another because patients with no expression values for the U6 or for pyknon-regions were excluded. Two-sided *t*-test has been used to evaluate differences between two groups. Y-axis values represent ratio of each pyknon-region to U6: ratios were calculated with the 2^−ΔCt^ method using U6 levels for normalization. **c,** Kaplan Meier curve reveals a poor clinical prognosis for patients with high pyk-reg-90 expression (*p*=0.016, log-rank test).

The above findings in cell lines and in paired normal/CRC samples indicated that the transcript corresponding to the pyk-reg-90 pyknon instance warranted further study. In what follows, and for reasons that will become evident shortly, we will be referring to the transcript captured by the pyk-reg-90 qRT-PCR assay as “N-BLR.”

### Cloning of the N-BLR lncRNA and *in situ* hybridization

The N-BLR transcript has its source in the 3p21.1-3p21.2 locus on the forward strand of chromosome 3. Targeted sequencing allowed us to zoom in on a 20 Kbp region from which we were able to clone N-BLR, an 845-nt transcript, originally from HCT116 and Colo320 cells as well as from normal colon. Subsequent Sanger sequencing confirmed that the same exact sequence, in terms of nucleotide content and length, was cloned from all three sources. This lncRNA is transcribed from the forward strand of chromosome 3 in the intergenic space between the WDR51A locus and the ALAS locus and corresponds to a contiguous block of genomic DNA (i.e. it is not spliced). WDR51A is located on the *reverse* strand, i.e. on the opposite strand from N-BLR and its transcription start site (TSS) is approximately 1.2 kb upstream from N-BLR (**Supp. Figure 3**). ALAS is on the same strand as N-BLR but more than 40 Kb downstream from it. Notably, N-BLR does not harbor any open reading frames (Hackanson et al. 2008) of significant length suggesting lack of protein-coding potential. Moreover, we verified that at the pyk-reg-90 locus transcription is preferentially derived from the forward strand i.e. it is sense to pyk-reg-90, and, thus to N-BLR (**Supp. Figure 4A**). We also screened for additional transcripts using primers targeting flanking regions 1, 2.5 and 5kb beyond N-BLR and in both genomic orientations: in nearly all cases, the qRT-PCR-identified transcripts were expressed at levels lower than those of N-BLR; the only exception was in the immediate 5′ region of N-BLR, where WDR51A gene is located (**Supp. Figure 4B**). Lastly, we performed *in situ* hybridization (ISH) in CRC and in adjacent normal colon samples, using a custom-designed locked-nucleic-acid probe: ISH reveals a 2–fold higher expression of N-BLR in tumor tissue versus matched normal colon sections (Fig. 2a), in agreement with qRT-PCR findings on N-BLR expression.

**Figure 2.**
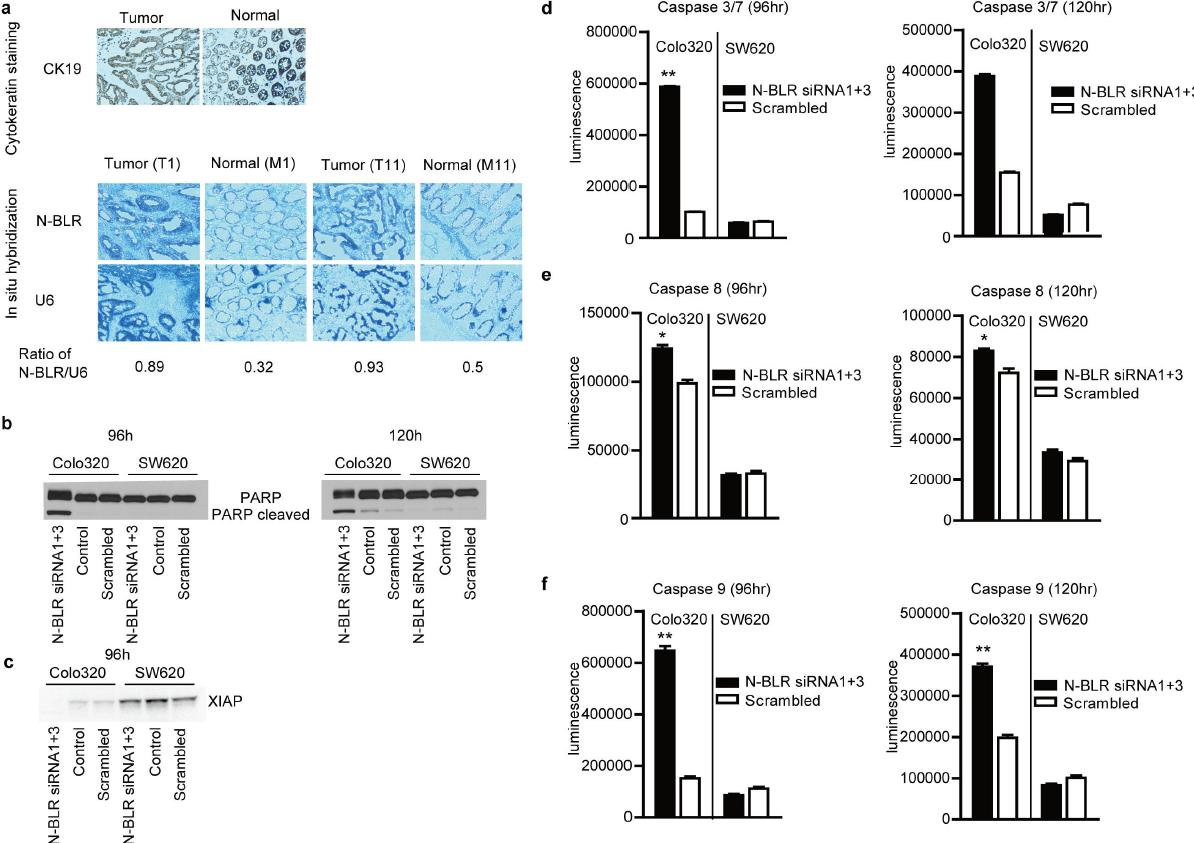
Properties of N-BLR. **a,** *In situ hybridization* shows differential expression of N-BLR in colon cancer and normal colon: hybridization for cytokeratin 19 (CK19) and RNA for N-BLR in colon cancer and in paired normal tissue showed epithelial expression and increased N-BLR expression in tumor comparing to paired normal tissue. **b,** PARP-1 expression following transfection of Colo320 and SW620 cells with siRNAs (N-BLR siRNA1+3 pool) against N-BLR. Profiling was carried out at 96 and 120 hrs. c: XIAP expression following transfection of Colo320 and SW620 cells with siRNAs (N-BLR siRNA1+3 pool) against N-BLR. **d, e, f:** expression of Casp3/7, Casp8 and Casp9 respectively following transfection of Colo320 and SW620 cells with siRNAs (N-BLR siRNA1+3 pool) against N-BLR. Profiling was carried out after 96 and 120 hrs.

### N-BLR is a novel component of the apoptotic pathway

We selected to experiment with Colo320 cells where N-BLR is up-regulated compared to SW620 cells (see **Supp. Figure 1**). The expression of N-BLR in SW620 is comparable to that in normal colon so we used SW620 as control. We designed four siRNAs (labeled as N-BLR siRNA1, siRNA2, siRNA3 and siRNA4) each of which efficiently targeted N-BLR. We combined two of the siRNAs (N-BLR siRNA1 and siRNA3) in a siRNA pool that reduced the clone's expression to less than 30%, in a dose dependent manner (**Supp. Figure 5A and 5B**). A non-targeting siRNA comprising a random sequence was used as a control. After treatment with the siRNA pool, N-BLR expression began decreasing at 24 hrs reaching minimum at 96 hrs in both cell lines. Cell counts of Colo320, but not of SW620, were significantly decreased at 96 hrs after treatment with either one of the single N-BLR siRNA1 and siRNA3 or the N-BLR siRNA1+3 pool (**Supp. Figure 5C**).

Apoptotic profiling of Colo320 cells at 96 and 120 hrs revealed significantly increased expression of cleaved PARP-1, a substrate of Caspase-3 and Caspase-7 compared to control following siRNA treatment with the N-BLR siRNA1+3 pool (Fig. 2b). Expression of the X-linked inhibitor of apoptosis (XIAP), an inhibitor of cell-death proteases Caspase-3 and Caspase-7, was abolished in Colo320 cells treated with siRNAs against N-BLR (p<0.001), but not in SW620 cells (Fig. 2c). The levels of Caspase-3, Caspase-7, Caspase-8, and Caspase-9 were significantly increased after siRNA targeting of N-BLR in Colo320 cells, but not in SW620 (Fig. 2d, 2e and 2f). The higher apoptosis in Colo320, but not SW620, was further confirmed by cell cycle analyses (**Supp. Figure 5D** and **5E**). Thus, N-BLR is a previously unknown component of the apoptotic pathway.

### N-BLR promotes invasion and migration

By transient down-regulation of N-BLR with two siRNAs, siRNA1+3 pool (**Supp. Figure 6A)**, we found that the number of migrated HCT116 cells in a transwell assay is significantly lower when compared to control (30-40% decrease – see **Supp. Figure 6B**). Similarly, we found that transient transfection with two other siRNAs, N-BLR siRNA3 and siRNA4 – S**upp. Figure 6C**, reduced the invasion potential of HCT116 cells (**Supp. Figure 6D** showing E-cadherin expression). To investigate further the effect of N-BLR down-regulation during tumorigenesis, we established *stable* N-BLR shRNA expressing clones (labeled as clone #3-1 and clone #4-7) and used them with HCT116 cells – the qRT-PCR assays indicate that the HCT116 cells are concordant with Colo320 cells with regard to the expression of the corresponding genomic transcripts – where they significantly inhibited the expression of N-BLR (Fig. 3a). We independently tested these clones versus stable clones that carried an empty vector in transwell based motility assays and found that down-regulation of N-BLR led to a concomitant decrease by more than 50% in *invasion* and an over 60% reduction in *migration* (Fig. 3b and 3c). Following transfection with each stable clone (vs. stable transfection with empty vectors) we carried out microarray-based analysis of gene expression: we found E-cadherin to be among the most up-regulated genes, and Vimentin among the most down-regulated ones (Fig. 3d). This is notable since E-cadherin and Vimentin are involved in the EMT and cell motility control in human colon carcinoma (Greco et al. 2010). We confirmed these findings by qRT-PCR (Fig. 3e), immunofluorescence (Fig. 3f) and Western blotting (Fig. 3g). Furthermore, the down-regulation of Vimentin was paralleled by down-regulation of ZEB1 (Fig. 3g and 3h), the latter being a known master regulator of EMT in human cancers.

**Figure 3.**
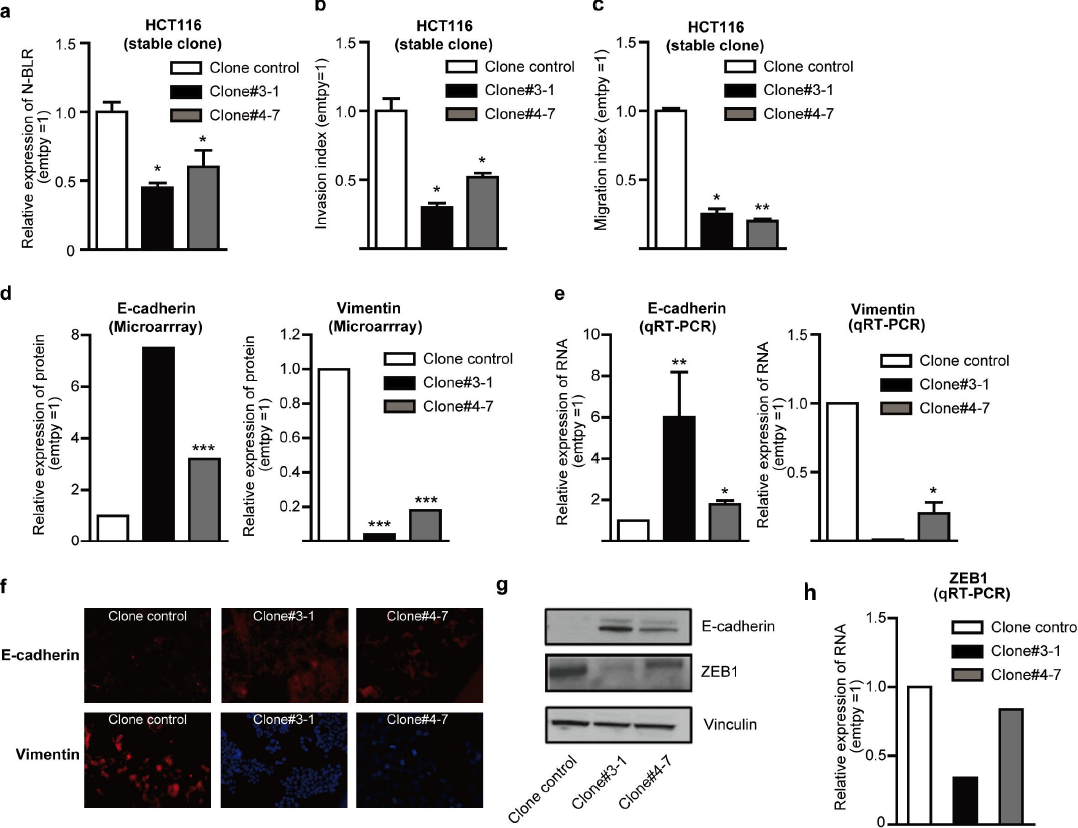
The effects on invasion of N-BLR down-regulation by specific siRNAs. **a**, N-BLR expression in stable clones. N-BLR expression is decreased in stably silenced clones. **b**, Invasion assays show significant reduction after 48 hrs for cells with stably silenced N-BLR. **c**, Migration assay at 24 hrs identified also significant reduction in migration of N-BLR stably silenced clones. **d**, Significantly differential expression of E-cadherin and vimentin identified by 44K Agilent microarray in HCT116 stable shRNA N-BLR clones #3-1 and #4-7 compared to control HCT 116 empty vector clone. **e**, Confirmation of microarray data by using real time PCR shows that E-cadherin is increased and vimentin is markedly decreased in both clones (#3-1 and #4-7). **f**, E-cadherin and vimentin were identified *in vivo* by immunofluorescence with specific antibodies. Immunofluorescence signal of E-cadherin (red color) was markedly increased in both clones. The vimentin signal was present in cells with Empty vector (red color) but not in clones #3-1 and #4-7. Blue color dots indicates nucleus. **g**, Western blotting for vinculin (loading control), E-cadherin and ZEB1 measured in the same clones. **h**, ZEB1 mRNA down-regulation in HCT116 stable shRNA N-BLR clones #3-1 and #4-7 compared to control HCT 116 empty vector clone.

### N-BLR is regulated by endogenous miRNAs

We next examined the possibility that N-BLR is regulated by miRNAs. In particular, we focused on the miR-200 family of miRNAs, a family with known involvement in the EMT (Gregory et al. 2008; Song et al. 2013). Of the five family members two, miR-141 and miR-200c, are endogenous and over-expressed after transient down-regulation of N-BLR in Colo320 cells (Fig. 4a). MiR-200a is expressed at low levels and increased only in clone#3-1 but not in clone #4-7 (data not shown) and therefore not further considered for the study. These two miRNAs are also significantly over-expressed in the stable clones #3-1 and #4-7 (Fig. 4b). Furthermore, the levels of N-BLR were decreased about 70% in HCT116 cells after miR-141 transfection and about 40% after miR-200c transfection (**Supp. Figure 7**). To determine whether the interaction between the two miRNAs and N-BLR is direct we used the RNA22 algorithm (Miranda et al. 2006) to look for target sites in the full 845 nt sequence of N-BLR. The algorithm predicted several potential binding sites for miR-141 and miR-200c (Fig. 4c), which were confirmed by luciferase assays (**Supp Fig. 8**). We also measured the expression of miR-141 in CRC samples that over-expressed N-BLR and found a significantly lower expression compared to matched normal tissue samples (Fig. 4d). Finally, using ISH we identified miR-200c in matched tumor and normal samples and measured an ∼50% reduction of expression in tumors versus normal, and an epithelium-specific expression (Fig. 4e).

**Figure 4.**
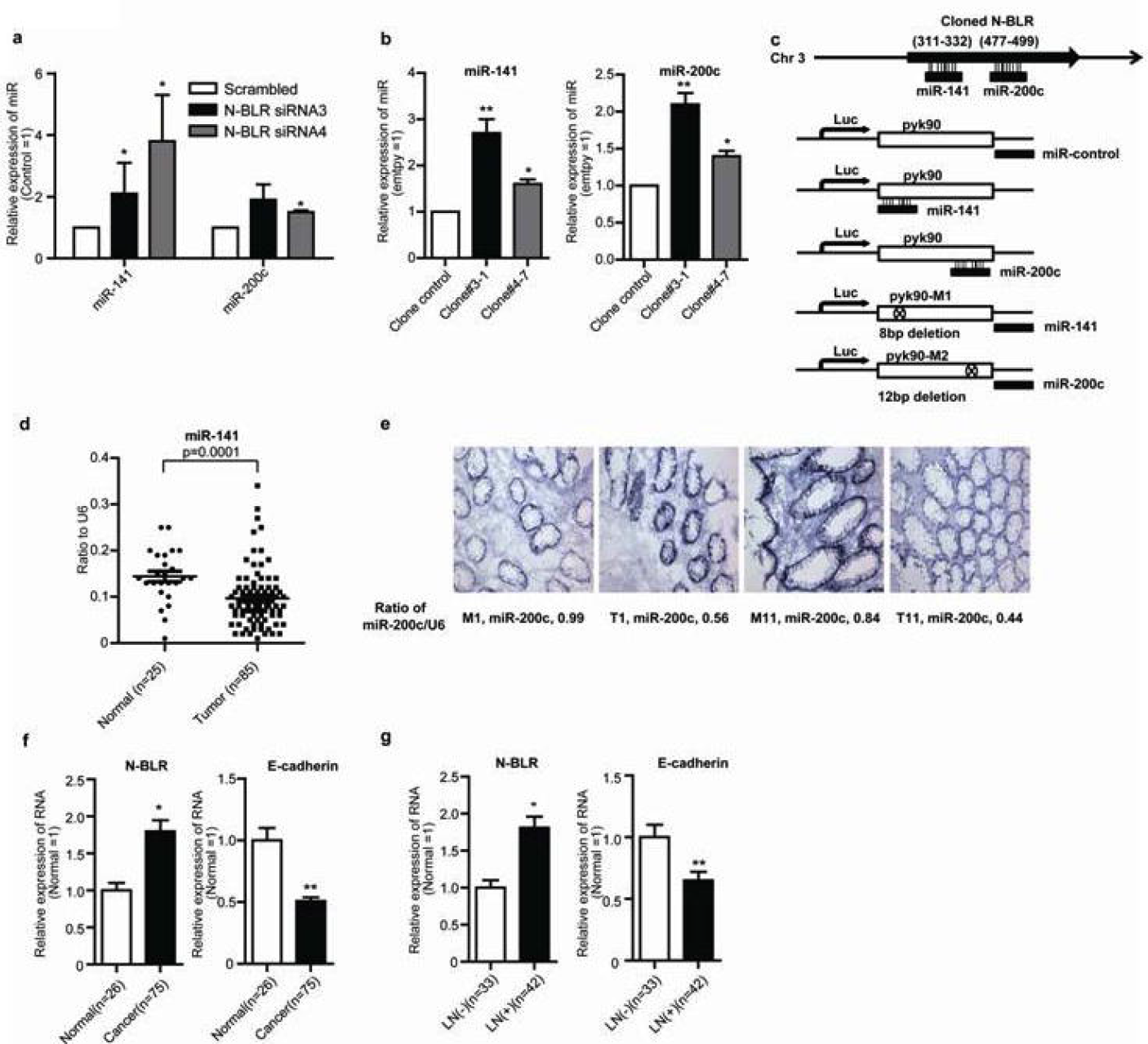
Interaction between N-BLR and miR-200 family members. **a**, The effect of transient transfection of N-BLR siRNA3 and siRNA4 on the miR-200 family in Colo320 cells. MiR-141 and miR-200c are increased in both N-BLR siRNAs transfected cells compared to scrambled control. **b,** miRNA expression in stable shRNA clones. MiR-141 and miR-200c are increased in both clones (#3-1 and #4-7). **c,** Schematic of the location of interaction sites between N-BLR and the miRNAs. **d,** miR-141 was down-regulated in CRC samples with respect to normal controls when measured by qRT-PCR. **e,** *In situ* hybridization for miR-200c in colon cancer and paired normal tissue. CK19 showed epithelial expression and U6 normalizer expression are as in Fig. 2. **f,** N-BLR and E-cadherin expression according to tumor and lymph node metastasis. N-BLR is increased and E-cadherin is decreased in CRC when compared to normal colon. **g,** The same is true when CRC with lymph node invasion (LN+) were compared with cases without lymph node involvement (LN-). Asterisks mark cases with statistically significant difference compared with scrambled (*p≤0.05; **p≤0.01).

### N-BLR expression modulates the EMT

Recent work revealed a novel modality in molecular interactions where two or more transcripts ‘regulate’ one another by competing for binding to the same miRNA (or group of miRNAs); the participating transcripts are called “competing endogenous RNAs” or *ceRNAs* (Poliseno et al. 2010; Rigoutsos and Furnari 2010; Cesana et al. 2011; Karreth et al. 2011; Sumazin et al. 2011; Tay et al. 2011). As mentioned, the miR-200 family has been linked to the EMT: indeed its members target, among other genes, the ZEB1/ZEB2 transcription factors that are repressors of E-cadherin expression (Gregory et al. 2008). At the same time, we have shown that two of the family's members, miR-141 and miR-200c, target and down-regulate N-BLR, which in turn has a inverse relationship with E-cadherin. Importantly, the inverse correlation between the expressions of N-BLR and E-cadherin extends to and has been observed in our cohort of CRC patients (Fig. 4f). The inverse correlation continues to hold when we compared the lymph-node positive (i.e. metastases to the lymph nodes) versus negative tumors (Fig. 4g).

Together, these findings support a model whereby N-BLR decoys ZEB1 by competing with them for binding to miR-141 and miR-200c. Consequently, an increase in the expression of N-BLR as the one we observe in the cell lines and in the CRC samples will attract the attention of a concomitantly higher number of copies from the available endogenous miR-141/miR-200c relieving pressure from and thereby up-regulating ZEB1. This will lead to a subsequent decrease of E-cadherin conferring the cells a mesenchymal phenotype that is correlated with increased invasiveness and migratory potential.

### Genome-wide profiling of pyknon transcripts

In light of the above many and diverse observations, we conjectured that the instances of the pyknon DNA motifs might serve as ‘homing beacons’ of sorts for genomic loci from which non-coding transcripts, potentially with functional relevance, could be transcribed. We set out to investigate this possibility by designing a custom microarray. There are more than 209,000 pyknons in the human genome (Rigoutsos et al. 2006; Tsirigos and Rigoutsos 2008) that have, by design, a minimum of 30 <u>intergenic/intronic</u> instances (=exact copies) and at least one more <u>exonic</u> instance. A standardized list of all known human pyknons together with a complete list of their coordinates across the span of the human genome is available at http://cm.jefferson.edu/pyknons.html. Among these more than 6 million instances, we randomly sub-selected a little over one thousand locations for our microarray by focusing on intergenic and intronic (i.e. non-protein-coding) instances that occur in the previously reported cancer associated genomic regions (Calin et al. 2004) (CAGRs). There are 1,292 such locations that correspond to 300 pyknons and are distributed across all chromosomes (**Supp. Figure 9**).

At each of the selected 1,292 locations, we *centered* a window of 100 nts and designed a 40-nt probe within it that overlapped with the corresponding pyknon; within the same window, we also designed a second 40-nt probe on the opposite strand that overlapped with the pyknon's *reverse complement*. This choice was dictated by our earlier published observation that for several thousand pyknons their reverse complement is also a pyknon (Rigoutsos et al. 2006). Of the 1,292 chosen locations 115 did not satisfy the probe design quality control criteria. The remaining 1,177 locations are probed in both the forward and reverse orientations for a grand total of 2,354 array probes. The microarray was augmented to also include probes for all known human miRNAs.

The 40-nt probes that we used to construct the array fall into two categories: those that have unique instances on the genome and those that have multiple instances. It is worth emphasizing that both the unique and non-unique array probes are important. Indeed, the incorporation of both types was by design. Each one of the unique probes allows to unambiguously interrogate a *single* location and to determine whether there is a transcriptional product from that region in one or more of the examined tissues. On the other hand, non-unique probes also have merit as they permit us to interrogate *multiple* genomic locations simultaneously, at the expense of losing localization information. Inclusion of non-unique probes was meant to help provide information along the lines of recent findings whereby protein-coding and non-protein-coding transcripts can modulate without direct molecular contact by competing for the same miRNA(s) that target them. Each such transcript can be thought of “decoying” the remaining transcript with which it shares a miRNA-targeting site.

### Unique and non-unique intergenic probes reveal tissue-specific expression profiles

We collected 15 normal samples from different individuals that spanned nine different tissues (4 colon, 2 breast, 1 lung, 1 heart, 1 skeletal muscle, 1 testicle, 1 liver, 2 mononuclear cells and 2 B-lymphocytes). We used our microarray to examine potential expression detected by the probes while ensuring that we distinguished among intergenic unique probes, intergenic non-unique probes, intronic unique probes and intronic non-unique probes. For the purposes of investigating the possibility of tissue-specific expression we focused solely on intergenic unique probes: we found several pyknon profiles that cluster according to the tissue of origin, which in turn suggests the existence of tissue-specific pyknon signatures (Fig. 5a). Not surprisingly, extending the analysis to include non-unique intergenic probes revealed similar tissue-specific expression (data not shown). This indicates that a sizable and wide-ranging spectrum of ceRNAs that compete for the attention of the same molecule, e.g. miRNA (Poliseno et al. 2010; Karreth et al. 2011; Tay et al. 2011; Ala et al. 2013; Song et al. 2013), DNMT1 (Di Ruscio et al. 2013), etc. is at play in different tissues. By using the same set of non-unique intergenic probes, compared to miRNAs, the pyknon-region transcripts exhibit higher tissue specificity in normal tissues as gauged by the Spearman correlation (**Supp. Figure 10**). Moreover, across all studied normal samples, the pyknon-regions exhibit significantly higher expression levels than miRNAs: **Supp. Figure 11A** shows the probability density functions of the normalized miRNA and pyknon probe intensities. The higher expression levels for the pyknon-regions were also observed in multiple additional normal samples (normal colon, B-lymphocytes, and mononuclear cells) from independent individuals (data not shown). Using an independent modality (qRT-PCR), we confirmed the data obtained from the array for select pyknon-regions comparing leukemia samples with normal B cell counterparts (see for example pyk-reg-14 in **Supp. Figure 11B**).

**Figure 5.**
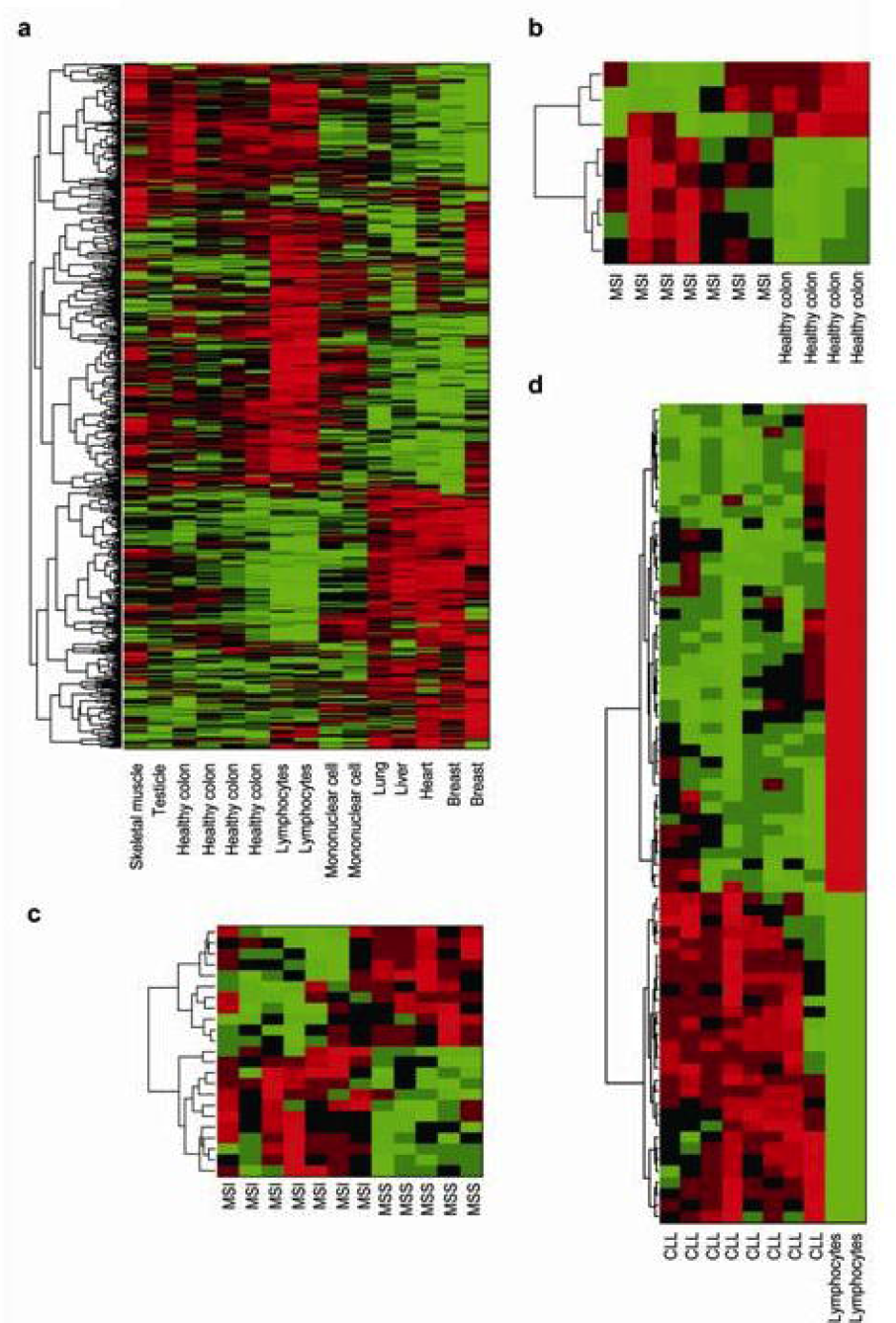
**a,** Pyknon clusters showing tissue specificity: the clusters were formed by considering solely unique intergenic probes. **b,** An example of a pyknon signature formed solely of intergenic unique probes that can distinguish healthy colon samples from microsatellite-unstable (MSI) samples. **c,** An example of a pyknon signature formed solely of intergenic unique probes that can distinguish MSI from microsatellite-stable (MSS) samples. **d,** A pyknon signature formed solely of intergenic unique probes that can distinguish CLL samples from healthy B-cell samples.

### Unique intergenic probes reveal differential expression between normal and disease samples

We also used our array to examine potential expression of the pyknons in *disease* samples. In Fig. 5b we show an example pyknon signature comprising unique intergenic probes that can distinguish healthy colon samples from microsatellite-unstable (MSI) samples. Interestingly, a subset of unique intergenic probes can correctly distinguish the two main categories of CRC, the microsatellite stable (MSS) and the MSI among the tested samples (Fig. 5c).

In addition to CRC, which is relevant for the work we presented above, we also examined CLL and normal B-lymphocyte samples, in order to explore the potential that our findings extend to other disease contexts. As with the normal samples, the expression profiles of the unique intergenic probes clustered according to the tissue of origin. Fig. 5d shows an example pyknon signature that can distinguish CLL samples from normal B-lymphocytes.

## Discussion

In this work, we presented our findings on N-BLR, a non-coding RNA that is a novel member of the apoptotic pathway where it acts as an inhibitor of apoptosis. In particular, we were able to show that inhibition of N-BLR led to down-regulation of XIAP and a subsequent up-regulation of the cell-death proteases Caspase-3 and Caspase-7, both of which are inhibited by XIAP. This suggests that those CRC settings where N-BLR is up-regulated are also associated with an increased resistance to apoptosis.

Additionally, N-BLR is able to modulate the EMT. This modulation is achieved mechanistically through N-BLR's direct interactions with at least two miRNAs (miR-141 and miR-200c) and indirect interactions with at least three mRNAs (ZEB1, E-cadherin, Vimentin). These six transcripts (N-BLR, miR-141, miR-200c, ZEB1, E-cadherin, Vimentin) represent a new signaling pathway in EMT. Our findings are supported by a recent article reporting that low expression of miR-200c on the invasive front of the primary CRCs and overexpression of miR-200c in CRC cells result in increased E-cadherin and reduced Vimentin expression (Hur et al. 2012). As we were preparing to submit our findings, another group reported analogous results for a different lncRNA in a different disease context (Yuan et al, 2014). In particular, they reported on an lncRNA, lncRNA-ATB, that promotes invasion and metastasis in hepatocellular carcinoma through interactions with members of the miR-200 family and with ZEB1/ZEB2. LncRNA-ATB is located on the reverse strand of chr14 between location 19858732 and 19861177 (hg19 assembly) and is a composite of three LINE-1 retrotransposon fragments and one full-length SINE retrotransposon. Each of these four component-fragments appears at numerous other locations on the human genome which in turn raises the possibility that very complex interactions such as those that we described in (Rigoutsos et al. 2006; Tsirigos and Rigoutsos 2008) and more recently in (Rigoutsos 2010) may be at play.

The second half of our presentation focuses on this very last point. Recall that what made us embark on the work with N-BLR was our earlier findings of a category of DNA motifs, the pyknons, together with a hypothesis that pyknon instances may pinpoint the locations of non-coding transcripts worth exploring. Indeed, the very sequence of N-BLR contains several instances of pyknon motifs. To pursue our hypothesis and investigate whether other pyknon loci on the genome show evidence of transcription we designed and built a custom microarray. The array comprised two sets of 40-nt probes: probes that were unique to the human genome, and probes that had multiple genomic instances. The rationale for the first category of probes is clear. Our choice of non-unique probes was informed by recent findings whereby protein-coding and non-protein-coding transcripts that are targeted by the same miRNA(s) can modulate one another in the absence of direct molecular interaction, a process that has been alternatively known as “decoying,” “quenching,” or “sponging” (Ebert and Sharp 2010b; Ebert and Sharp 2010a; Poliseno et al. 2010; Karreth et al. 2011; Tay et al. 2011; Ala et al. 2013; Song et al. 2013). Even though the second group of probes cannot, without additional effort, uniquely identify the genomic source of the ncRNA(s) hybridizing to them, it does demonstrate that at least one and perhaps multiple such transcripts exist. This finding and the fact that the corresponding underlying pyknon motif is identically present in at least one mRNA (this being the very definition of pyknon motifs) raise the distinct possibility that these ncRNAs can act as molecular decoys for the corresponding mRNA. With regard to the second goal, our analyses of unique intergenic probes revealed several pyknon-containing regions whose expression profiles are tissue-and cell-state (disease/normal) dependent. This in turn indicates that there are several non-coding transcripts that are involved in specific contexts and which currently remain uncharacterized.

In closing, it is worth emphasizing that additional examination of the N-BLR locus revealed that it is primate-specific and thus not conserved in rodents. As such, N-BLR's activity cannot be captured by mouse models of colon cancer. This represents another intriguing dimension of the intricacies of human disease and of N-BLR's regulatory activity of the EMT. In that regard, N-BLR and other similar molecules are unlike miRNAs (Esquela-Kerscher and Slack 2006; Hammond 2007; Mendell 2008; Slack and Weidhaas 2008), transcribed UCRs (Calin et al. 2007b; Scaruffi et al. 2009; Mestdagh et al. 2010), and lincRNAs (Gupta et al. 2010b). Organism-specific transcripts can be thought of as representing a paradigm shift that is in agreement with the increasing realization that human cancers differ from animal models involving the same gene and the specific human mutation (Van Dyke and Jacks 2002). Naturally, their sequence properties make N-BLR and molecules like it promising as novel prognostic indicators.

## METHODS

Methods and any associated references are available in the online version of the paper.

## Acknowledgements

We thank Dr. Stanley Hamilton Department of Pathology, The University of Texas M. D. Anderson Cancer Center, Houston, TX, USA for the continuous support and critical reading of the manuscript. We thank Grace Zhang and Howard Chang for sharing with us the genomic locations of lincRNAs. The GeneBank accession numbers for the cloned pyknon-containing-regions described in this study are: HQ262399, HQ262400, HQ262401 and HQ262402. The Array Express submission number is E-MTAB-298. Dr. Yeh's work was supported by the NIH/NCI (CA140424). We thank the UNC PDX Program for their assistance. Dr Calin is The Alan M. Gewirtz Leukemia & Lymphoma Society Scholar. He was supported also as a Fellow at The University of Texas MD Anderson Research Trust, as a University of Texas System Regents Research Scholar and by the CLL Global Research Foundation. Work in Dr. Calin's laboratory is supported in part by the NIH/NCI grants 1 R01 CA182905-01 and 1UH2TR00943-01, T32CA009599, Developmental Research Awards in Prostate Cancer, Multiple Myeloma, Leukemia (P50 CA100632) and Head and Neck (P50 CA097007) SPOREs, a SINF MDACC_DKFZ grant in CLL, a SINF grant in colon cancer, a Kidney Cancer Pilot Project, a 2013 Sprint for Life Research Award the Laura and John Arnold Foundation, the RGK Foundation and the Estate of C. G. Johnson, Jr. Work in Dr. Rigoutsos’ laboratory was supported in part by a 2011 William M. Keck Foundation Medical Research Award, in part by institutional funds, and in part by a grant from the Pennsylvania Department of Health which specifically disclaims responsibility for any analyses, interpretations or conclusions.

## Author contributions

G.A.C. and I.R. designed the experiments. S.K.L, S.Y.N., T.C I., M.P., H.L., Y.J., M.Shi., R.S.R. and M.Y.S. performed most of the biological experiments including tissue processing, tissue culture experiments, qRT-PCR and investigation of the biological effects. I.R., S.R., P.C., A.T., L.X. and G.A.C. performed most of the computational experiments and analyzed the data. J.J.Y. provided samples. R.G., M.N., E.F., M.J.K. and G.L. provided samples and performed clinical data correlations. E.J.J, R.S., M.S.N., A.T., and R.T. performed tissue culture experiments and vectors construction. S.A.M., E.F., and M.E. provided key experimental reagents. X.Z. designed probes and performed the ISH experiments. G.A.C. and C.G.L. participated in the custom array design. G.A.C., I.R., S.K.L and S.Y.N. wrote the manuscript. All authors discussed the results and read and commented on the manuscript's contents during the initial submission and the subsequent revisions.

## REFERENCES

Ala U, Karreth FA, Bosia C, Pagnani A, Taulli R, Leopold V, Tay Y, Provero P, Zecchina R, Pandolfi PP. 2013. Integrated transcriptional and competitive endogenous RNA networks are cross-regulated in permissive molecular environments. Proceedings of the National Academy of Sciences of the United States of America.

Ambros V. 2008. The evolution of our thinking about microRNAs. Nature medicine 14(10): 1036–1040.

Bartel DP. 2004. MicroRNAs: genomics, biogenesis, mechanism, and function. Cell 116(2): 281–297.

Bartel DP. 2009. MicroRNAs: target recognition and regulatory functions. Cell 136(2): 215–233.

Bertone P, Stolc V, Royce TE, Rozowsky JS, Urban AE, Zhu X, Rinn JL, Tongprasit W, Samanta M, Weissman S et al. 2004. Global identification of human transcribed sequences with genome tiling arrays. Science (New York, NY 306(5705): 2242–2246.

Bischof JM, Stewart CL, Wevrick R. 2007. Inactivation of the mouse Magel2 gene results in growth abnormalities similar to Prader-Willi syndrome. Human Molecular Genetics 16(22): 2713–2719.

Calin GA, Liu C-G, Ferracin M, Hyslop T, Spizzo R, Sevignani C, Fabbri M, Cimmino A, Lee EJ, Wojcik SE et al. 2007a. Ultraconserved regions encoding ncRNAs are altered in human leukemias and carcinomas. Cancer Cell 12(3): 215–229.

Calin GA, Liu CG, Ferracin M, Hyslop T, Spizzo R, Sevignani C, Fabbri M, Cimmino A, Lee EJ, Wojcik SE et al. 2007b. Ultraconserved regions encoding ncRNAs are altered in human leukemias and carcinomas. Cancer Cell 12(3): 215–229.

Calin GA, Sevignani C, Dumitru CD, Hyslop T, Noch E, Yendamuri S, Shimizu M, Rattan S, Bullrich F, Negrini M et al. 2004. Human microRNA genes are frequently located at fragile sites and genomic regions involved in cancers. Proc Natl Acad Sci U S A 101(9): 2999–3004.

Carninci P, Kasukawa T, Katayama S, Gough J, Frith MC, Maeda N, Oyama R, Ravasi T, Lenhard B, Wells C. et al. 2005. The transcriptional landscape of the mammalian genome. Science (New York, NY 309(5740): 1559–1563.

Cesana M, Cacchiarelli D, Legnini I, Santini T, Sthandier O, Chinappi M, Tramontano A, Bozzoni I. 2011. A Long Noncoding RNA Controls Muscle Differentiation by Functioning as a Competing Endogenous RNA. Cell 147(2): 358–369.

Chen L-L, Carmichael GG. 2009. Altered nuclear retention of mRNAs containing inverted repeats in human embryonic stem cells: functional role of a nuclear noncoding RNA. Molecular Cell 35(4): 467–478.

Cheng J, Kapranov P, Drenkow J, Dike S, Brubaker S, Patel S, Long J, Stern D, Tammana H, Helt G et al. 2005. Transcriptional maps of 10 human chromosomes at 5-nucleotide resolution. Science (New York, NY 308(5725): 1149–1154.

Di Ruscio A, Ebralidze AK, Benoukraf T, Amabile G, Goff LA, Terragni J, Figueroa ME, De Figueiredo Pontes LL, Alberich-Jorda M, Zhang P et al. 2013. DNMT1-interacting RNAs block gene-specific DNA methylation. Nature: 1–10.

Djebali S, Davis CA, Merkel A, Dobin A, Lassmann T, Mortazavi A, Tanzer A, Lagarde J, Lin W, Schlesinger F et al. 2012. Landscape of transcription in human cells. Nature 489(7414): 101–108.

Ebert MS, Sharp PA. 2010a. Emerging Roles for Natural MicroRNA Sponges. Current Biology 20(19): R858–R861.

Ebert MS, Sharp PA. 2010b. MicroRNA sponges: Progress and possibilities. RNA (New York, NY) 16(11): 2043–2050.

Esquela-Kerscher A, Slack FJ. 2006. Oncomirs -microRNAs with a role in cancer. Nat Rev Cancer 6(4): 259–269.

Feng J, Naiman DQ, Cooper B. 2009. Coding DNA repeated throughout intergenic regions of the Arabidopsis thaliana genome: evolutionary footprints of RNA silencing. Mol Biosyst 5(12): 1679–1687.

Flynn RA, Zheng GXY, Raj A, Rinn JL, Chang HY. 2012. Control of somatic tissue differentiation by the long non-coding RNA TINCR. Nature.

Greco C, Bralet M, Ailane N, Dubart Kupperschmitt A, Rubinstein E, Le-Naour F, Boucheix C. 2010. E-cadherin/p120-catenin and tetraspanin Co-029 cooperate for cell motility control in human colon carcinoma. Cancer research 70(19): 7674–7683.

Gregory PA, Bert AG, Paterson EL, Barry SC, Tsykin A, Farshid G, Vadas MA, Khew-Goodall Y, Goodall GJ. 2008. The miR-200 family and miR-205 regulate epithelial to mesenchymal transition by targeting ZEB1 and SIP1. Nature Cell Biology 10(5): 593–601.

Gupta RA, Shah N, Wang KC, Kim J, Horlings HM, Wong DJ, Tsai M-C, Hung T, Argani P, Rinn JL et al. 2010a. Long non-coding RNA HOTAIR reprograms chromatin state to promote cancer metastasis. Nature 464(7291): 1071–1076.

Gupta RA, Shah N, Wang KC, Kim J, Horlings HM, Wong DJ, Tsai MC, Hung T, Argani P, Rinn JL et al. 2010b. Long non-coding RNA HOTAIR reprograms chromatin state to promote cancer metastasis. Nature 464(7291): 1071–1076.

Guttman M, Amit I, Garber M, French C, Lin MF, Feldser D, Huarte M, Zuk O, Carey BW, Cassady JP et al. 2009. Chromatin signature reveals over a thousand highly conserved large non-coding RNAs in mammals. Nature 458(7235): 223–227.

Hackanson B, Bennett KL, Brena RM, Jiang J, Claus R, Chen SS, Blagitko-Dorfs N, Maharry K, Whitman SP, Schmittgen TD et al. 2008. Epigenetic modification of CCAAT/enhancer binding protein alpha expression in acute myeloid leukemia. Cancer research 68(9): 3142–3151.

Hammond SM. 2007. MicroRNAs as tumor suppressors. Nature genetics 39(5): 582–583.

Hur K, Toiyama Y, Takahashi M, Balaguer F, Nagasaka T, Koike J, Hemmi H, Koi M, Boland CR, Goel A. 2012. MicroRNA-200c modulates epithelial-to-mesenchymal transition (EMT) in human colorectal cancer metastasis. Gut.

Hutchinson JN, Ensminger AW, Clemson CM, Lynch CR, Lawrence JB, Chess A. 2007. A screen for nuclear transcripts identifies two linked noncoding RNAs associated with SC35 splicing domains. BMC Genomics 8: 39.

Karreth FA, Tay Y, Perna D, Ala U, Tan SM, Rust AG, DeNicola G, Webster KA, Weiss D, Perez-Mancera PA et al. 2011. In Vivo Identification of Tumor-Suppressive PTEN ceRNAs in an Oncogenic BRAF-Induced Mouse Model of Melanoma. Cell 147(2): 382–395.

Khaitan D, Dinger ME, Mazar J, Crawford J, Smith MA, Mattick JS, Perera RJ. 2011. The Melanoma-Upregulated Long Noncoding RNA SPRY4-IT1 Modulates Apoptosis and Invasion. Cancer research 71(11): 3852–3862.

Khaitovich P, Kelso J, Franz H, Visagie J, Giger T, Joerchel S, Petzold E, Green RE, Lachmann M, Paabo S. 2006. Functionality of intergenic transcription: an evolutionary comparison. PLoS genetics 2(10): e171.

Kim VN. 2006. Small RNAs just got bigger: Piwi-interacting RNAs (piRNAs) in mammalian testes. Genes Dev 20(15): 1993–1997.

Kogo R, Shimamura T, Mimori K, Kawahara K, Imoto S, Sudo T, Tanaka F, Shibata K, Suzuki A, Komune S et al. 2011. Long Noncoding RNA HOTAIR Regulates Polycomb-Dependent Chromatin Modification and Is Associated with Poor Prognosis in Colorectal Cancers. Cancer research 71(20): 6320–6326.

Langenberger D, Bermudez-Santana C, Hertel J, Hoffmann S, Khaitovich P, Stadler PF. 2009. Evidence for human microRNA-offset RNAs in small RNA sequencing data. Bioinformatics (Oxford, England) 25(18): 2298–2301.

Latos PA, Pauler FM, Koerner MV, Senergin HB, Hudson QJ, Stocsits RR, Allhoff W, Stricker SH, Klement RM, Warczok KE et al. 2012. Airn Transcriptional Overlap, But Not Its lncRNA Products, Induces Imprinted Igf2r Silencing. Science (New York, NY) 338(6113): 1469–1472.

Lee JT, Lu N. 1999. Targeted mutagenesis of Tsix leads to nonrandom X inactivation. Cell 99(1): 47–57.

Louro R, Smirnova AS, Verjovski-Almeida S. 2009. Long intronic noncoding RNA transcription: expression noise or expression choice? Genomics 93(4): 291–298.

Maass PG, Rump A, Schulz H, Stricker S, Schulze L, Platzer K, Aydin A, Tinschert S, Goldring MB, Luft FC et al. 2012. A misplaced lncRNA causes brachydactyly in humans. J Clin Invest 122(11): 3990–4002.

Mattick JS. 2009. The genetic signatures of noncoding RNAs. PLoS genetics 5(4): e1000459.

Mattick JS, Makunin IV. 2006. Non-coding RNA. Human Molecular Genetics 15(Review Issue 1): R17.

Mendell JT. 2008. miRiad roles for the miR-17-92 cluster in development and disease. Cell 133(2): 217–222.

Mestdagh P, Fredlund E, Pattyn F, Rihani A, Van Maerken T, Vermeulen J, Kumps C, Menten B, De Preter K, Schramm A et al. 2010. An integrative genomics screen uncovers ncRNA T-UCR functions in neuroblastoma tumours. Oncogene 29(24): 3583–3592.

Miranda KC, Huynh T, Tay Y, Ang Y-S, Tam W-L, Thomson AM, Lim B, Rigoutsos I. 2006. A pattern-based method for the identification of MicroRNA binding sites and their corresponding heteroduplexes. Cell 126(6): 1203–1217.

Mukherji S, Ebert MS, Zheng GXY, Tsang JS, Sharp PA, van Oudenaarden A. 2011. MicroRNAs can generate thresholds in target gene expression. Nature genetics 43(9): 854–859.

Ogawa Y, Sun BK, Lee JT. 2008. Intersection of the RNA interference and X-inactivation pathways. Science (New York, NY) 320(5881): 1336–1341.

Orom UA, Derrien T, Beringer M, Gumireddy K, Gardini A, Bussotti G, Lai F, Zytnicki M, Notredame C, Huang Q et al. 2010. Long noncoding RNAs with enhancer-like function in human cells. Cell 143(1): 46–58.

Pandey RR, Mondal T, Mohammad F, Enroth S, Redrup L, Komorowski J, Nagano T, Mancini-Dinardo D, Kanduri C. 2008. Kcnq1ot1 antisense noncoding RNA mediates lineage-specific transcriptional silencing through chromatin-level regulation. Molecular Cell 32(2): 232–246.

Poliseno L, Salmena L, Zhang J, Carver B, Haveman WJ, Pandolfi PP. 2010. A coding-independent function of gene and pseudogene mRNAs regulates tumour biology. Nature 465(7301): 1033–1038.

Prensner JR, Iyer MK, Balbin OA, Dhanasekaran SM, Cao Q, Brenner JC, Laxman B, Asangani IA, Grasso CS, Kominsky HD et al. 2011. Transcriptome sequencing across a prostate cancer cohort identifies PCAT-1, an unannotated lincRNA implicated in disease progression. Nature Biotechnology 29(8): 744–758.

Rigoutsos I. 2009. New tricks for animal microRNAS: targeting of amino acid coding regions at conserved and nonconserved sites. Cancer research 69(8): 3245–3248.

Rigoutsos I. 2010. Short RNAs: how big is this iceberg? Curr Biol 20(3): R110–113.

Rigoutsos I, Floratos A. 1998. Combinatorial pattern discovery in biological sequences: The TEIRESIAS algorithm. Bioinformatics (Oxford, England) 14(1): 55–67.

Rigoutsos I, Furnari F. 2010. Gene-expression forum: Decoy for microRNAs. Nature 465(7301): 1016–1017.

Rigoutsos I, Huynh T, Miranda K, Tsirigos A, McHardy A, Platt D. 2006. Short blocks from the noncoding parts of the human genome have instances within nearly all known genes and relate to biological processes. Proc Natl Acad Sci U S A 103(17): 6605–6610.

Robine N, Lau NC, Balla S, Jin Z, Okamura K, Kuramochi-Miyagawa S, Blower MD, Lai EC. 2009. A broadly conserved pathway generates 3'UTR-directed primary piRNAs. Curr Biol 19(24): 2066–2076.

Saito K, Inagaki S, Mituyama T, Kawamura Y, Ono Y, Sakota E, Kotani H, Asai K, Siomi H, Siomi MC. 2009. A regulatory circuit for piwi by the large Maf gene traffic jam in Drosophila. Nature 461(7268): 1296–1299.

Scaruffi P, Stigliani S, Moretti S, Coco S, De Vecchi C, Valdora F, Garaventa A, Bonassi S, Tonini GP. 2009. Transcribed-Ultra Conserved Region expression is associated with outcome in high-risk neuroblastoma. BMC cancer 9: 441.

Slack FJ, Weidhaas JB. 2008. MicroRNA in cancer prognosis. The New England journal of medicine 359(25): 2720–2722.

Small EM, Olson EN. 2011. Pervasive roles of microRNAs in cardiovascular biology. Nature 469(7330): 336–342.

Song SJ, Poliseno L, Song MS, Ala U, Webster K, Ng C, Beringer G, Brikbak NJ, Yuan X, Cantley LC et al. 2013. MicroRNA-Antagonism Regulates Breast Cancer Stemness and Metastasis via TET-Family-Dependent Chromatin Remodeling. Cell: 1–14.

Sumazin P, Yang X, Chiu H-S, Chung W-J, Iyer A, Llobet-Navas D, Rajbhandari P, Bansal M, Guarnieri P, Silva J et al. 2011. An Extensive MicroRNA-Mediated Network of RNA-RNA Interactions Regulates Established Oncogenic Pathways in Glioblastoma. Cell 147(2): 370–381.

Taft RJ, Glazov EA, Cloonan N, Simons C, Stephen S, Faulkner GJ, Lassmann T, Forrest AR, Grimmond SM, Schroder K et al. 2009. Tiny RNAs associated with transcription start sites in animals. Nature genetics 41(5): 572–578.

Tay Y, Kats L, Salmena L, Weiss D, Tan SM, Ala U, Karreth F, Poliseno L, Provero P, Di Cunto F et al. 2011. Coding-Independent Regulation of the Tumor Suppressor PTEN by Competing Endogenous mRNAs. Cell 147(2): 344–357.

Tian D, Sun S, Lee JT. 2010. The Long Noncoding RNA, Jpx, Is a Molecular Switch for X Chromosome Inactivation. Seminars in Fetal and Neonatal Medicine 143(3): 390–403.

Tsirigos A, Rigoutsos I. 2008. Human and mouse introns are linked to the same processes and functions through each genome's most frequent non-conserved motifs. Nucleic acids research 36(10): 3484–3493.

Van Dyke T, Jacks T. 2002. Cancer modeling in the modern era: progress and challenges. Cell 108(2): 135–144.

Vasilescu C, Rossi S, Shimizu M, Tudor S, Veronese A, Ferracin M, Nicoloso MS, Barbarotto E, Popa M, Stanciulea O et al. 2009. MicroRNA fingerprints identify miR-150 as a plasma prognostic marker in patients with sepsis. PLoS ONE 4(10): e7405.

Wang KC, Yang YW, Liu B, Sanyal A, Corces-Zimmerman R, Chen Y, Lajoie BR, Protacio A, Flynn RA, Gupta RA et al. 2011. A long noncoding RNA maintains active chromatin to coordinate homeotic gene expression. Nature 472(7341): 120–124.

Williams AH, Liu N, van Rooij E, Olson EN. 2009. MicroRNA control of muscle development and disease. Current opinion in cell biology 21(3): 461–469.

Yang L, Lin C, Liu W, Zhang J, Ohgi KA, Grinstein JD, Dorrestein PC, Rosenfeld MG. 2011. ncRNA-and Pc2 methylation-dependent gene relocation between nuclear structures mediates gene activation programs. Cell 147(4): 773–788.

Yuan JH, Yang F, Wang F, Ma JZ, Guo YJ, Tao QF, Liu F, Pan W, Wang TT, Zhou CC, Wang SB, Wang YZ, Yang Y, Yang N, Zhou WP, Yang GS, Sun SH. A Long Noncoding RNA Activated by TGF-β Promotes the Invasion-Metastasis Cascade in Hepatocellular Carcinoma. Cancer Cell. 2014 Apr 22. pii: S1535-6108(14)00119-6. doi: 10.1016/j.ccr.2014.03.010. [Epub ahead of print]

Zhao J, Sun BK, Erwin JA, Song JJ, Lee JT. 2008. Polycomb Proteins Targeted by a Short Repeat RNA to the Mouse X Chromosome. Science (New York, NY) 322(5902): 750–756.

